# Inferring therapeutic vulnerability within tumors through integration of pan-cancer cell line and single-cell transcriptomic profiles

**DOI:** 10.1101/2023.10.29.564598

**Authors:** Weijie Zhang, Danielle Maeser, Adam Lee, Yingbo Huang, Robert F. Gruener, Israa G. Abdelbar, Sampreeti Jena, Anand G. Patel, R. Stephanie Huang

## Abstract

Single-cell RNA sequencing greatly advanced our understanding of intratumoral heterogeneity through identifying tumor subpopulations with distinct biologies. However, translating biological differences into treatment strategies is challenging, as we still lack tools to facilitate efficient drug discovery that tackles heterogeneous tumors. One key component of such approaches tackles accurate prediction of drug response at the single-cell level to offer therapeutic options to specific cell subpopulations. Here, we present a transparent computational framework (nicknamed scIDUC) to predict therapeutic efficacies on an individual-cell basis by integrating single-cell transcriptomic profiles with large, data-rich pan-cancer cell line screening datasets. Our method achieves high accuracy, with predicted sensitivities easily able to separate cells into their true cellular drug resistance status as measured by effect size (Cohen’s d > 1.0). More importantly, we examine our method’s utility with three distinct prospective tests covering different diseases (rhabdomyosarcoma, pancreatic ductal adenocarcinoma, and castration-resistant prostate cancer), and in each our predicted results are accurate and mirrored biological expectations. In the first two, we identified drugs for cell subpopulations that are resistant to standard-of-care (SOC) therapies due to intrinsic resistance or effects of tumor microenvironments. Our results showed high consistency with experimental findings from the original studies. In the third test, we generated SOC therapy resistant cell lines, used scIDUC to identify efficacious drugs for the resistant line, and validated the predictions with in-vitro experiments. Together, scIDUC quickly translates scRNA-seq data into drug response for individual cells, displaying the potential as a first-line tool for nuanced and heterogeneity-aware drug discovery.

## Introduction

Heterogeneity within tumors, where distinct cell subpopulations display varying features, has been causally linked to therapy resistance and disease recurrence in many cancers^1–3^. Tumor cells unresponsive to standard-of-care (SOC) pharmacological interventions continue to proliferate and cascade disease progression under the selective pressure. Such phenotypic aberrations often correlate with molecular variations in cellular mutational and transcriptional profiles^4–6^. Thanks to the quickly evolving single-cell (SC) sequencing technologies, genomic and transcriptomic landscapes within tumors in many patient populations have been continuously characterized^7,8^. On the other hand, the increasingly available single-cell sequencing data confers opportunities for development of new treatment strategies that tackle problematic cell groups, address clonal heterogeneity, and eventually help achieve curability in cancers^9,10^.

In addition to traditional pharmaceutical research and development pipelines, computational frameworks have emerged as an indispensable tool for drug discovery for their cost-efficient nature and more importantly, the ability to screen many drugs for various indications^11–13^. Current *in silico* drug discovery models are largely constructed based on openly available high-throughput drug screens on pan-cancer cell lines (CCLs)^14,15^, whose transcriptomic profiles are systematically evaluated through bulk RNA sequencing (RNA-seq)^16^. While computational tools utilizing relationships between CCL gene expression from RNA-seq and drug response have demonstrated practicality in predicting efficacious treatments^13,17–19^, such relationships cannot be directly applied to generate predictions of drug response at the cellular level, as RNA-seq is limited to measuring average expression across a diverse set of cells, which obscures cell type and composition, as well as temporal and spatial distributions. Thus, inferring cellular drug response requires specialized tools to transfer current bulk-learned drug-gene information to single-cell RNA sequencing (scRNA-seq) data that encapsulate cell level expression patterns^20–24^.

In recent years, such computational tools have been conceptualized, and a few implementations have also been proposed^10,20,24^. The common crucial functionality among the proposed methods relates to overcoming fundamental differences in properties between bulk and SC RNA-seq data to enable predictions of drug response at the SC resolution using learned drug-gene information from bulk data. To achieve this, Beyondcell, DREEP, and scDr choose to learn a fixed number of drug-specific signature genes from bulk CCL data and apply learned signatures independently in scRNA-seq data to calculate signature scores which indicate drug response^22,25,26^. However, considering that genes may harbor varying predictability for response to different drugs and scRNA-seq data are notorious for their low detection rates as well as stochastic drop-outs^14,27^, it is not guaranteed that gene signatures always deliver reliable predictions of sensitivities to various drugs^28^. In comparison, SCAD and scDEAL directly tackle differences between bulk and SC data and emphasize integration of the two domains via neural network based approaches^23,29^. While data-hungry deep learning (DL) routes could benefit from large scRNA-seq data and model complex drug-gene relationships, the availability of CCL bulk data could pose continued limits against accurate parameter estimation. Also, it has been shown that DL methods offer comparable performances as classical machine learning frameworks in drug response modeling^19^, while in general consisting of more parameters and request more computing resources. CaDRReS-Sc also conducts bulk-SC data integration but through projecting original data into a fixed-dimensional subspace^21^. These integration-embedded methods by default use dichotomous labels for drug response (sensitive or resistant); such arbitrary cutoffs may not reflect pharmacological properties and could mask variation of drug response among heterogeneous cells. Furthermore, insufficient evidence has been presented thus far to demonstrate the translational value of these predictive models in aiding cell-type aware early development with diverse biology models.

Therefore, to fill the current gap and to establish an adaptable virtual SC drug screen platform tailored toward clinically meaningful predictions, we present scIDUC (<u>s</u>ingle-<u>c</u>ell Integration and <u>D</u>r<u>U</u>g response <u>C</u>omputation), a novel and transparent transfer learning-based framework that quickly and accurately generates predictions of drug responses for scRNA-seq data. scIDUC learns relationships between drug sensitivities and relevant gene expression patterns based on CCL RNA-seq data and CCL high-throughput drug screens. Integration of CCL RNA-seq dataset and target scRNA-seq dataset is performed to denoise and extract shared gene expression patterns between bulk and SC data sources; the resulted bulk data is then used to train drug response models, whose coefficients are further applied to post-integration SC data to infer cellular drug sensitivity scores. We evaluated our method using a variety of scRNA-seq datasets with known cellular drug sensitivity status. Through prospective analysis in three distinct scenarios, we further demonstrated the versatility of our framework in various biological models addressing research questions and generating meaningful therapeutic predictions with potential clinical impact. Validation of predictions yielded from scIDUC substantiates its potential as a first-line research tool in the field of computational drug discovery for addressing intratumoral heterogeneity and to facilitate hypothesis formulation for various oncology research topics.

## Results

### Overall framework of scIDUC

An overview of our scIDUC computational pipeline is outlined in Figure 1. Drug screen results from the CTRPv2 in normalized area-under-the-dose-response-curve (nAUC) were used as CCL drug response (see Methods). Prior to main computation steps, we aimed to identify drug response relevant genes (DRGs, see Methods) for each drug (<u>Step 0</u>). Instead of filtering for a certain number of DRGs using an arbitrary threshold, a typical DRG list for a drug still consists of all genes, which are ranked from most drug response informative to the least. This is to partially address that different drug may display different predictability in different scRNA-seq data. The input bulk and SC RNA-seq datasets were then subset to retain the same DRGs to facilitate data integration.

**Figure 1.**
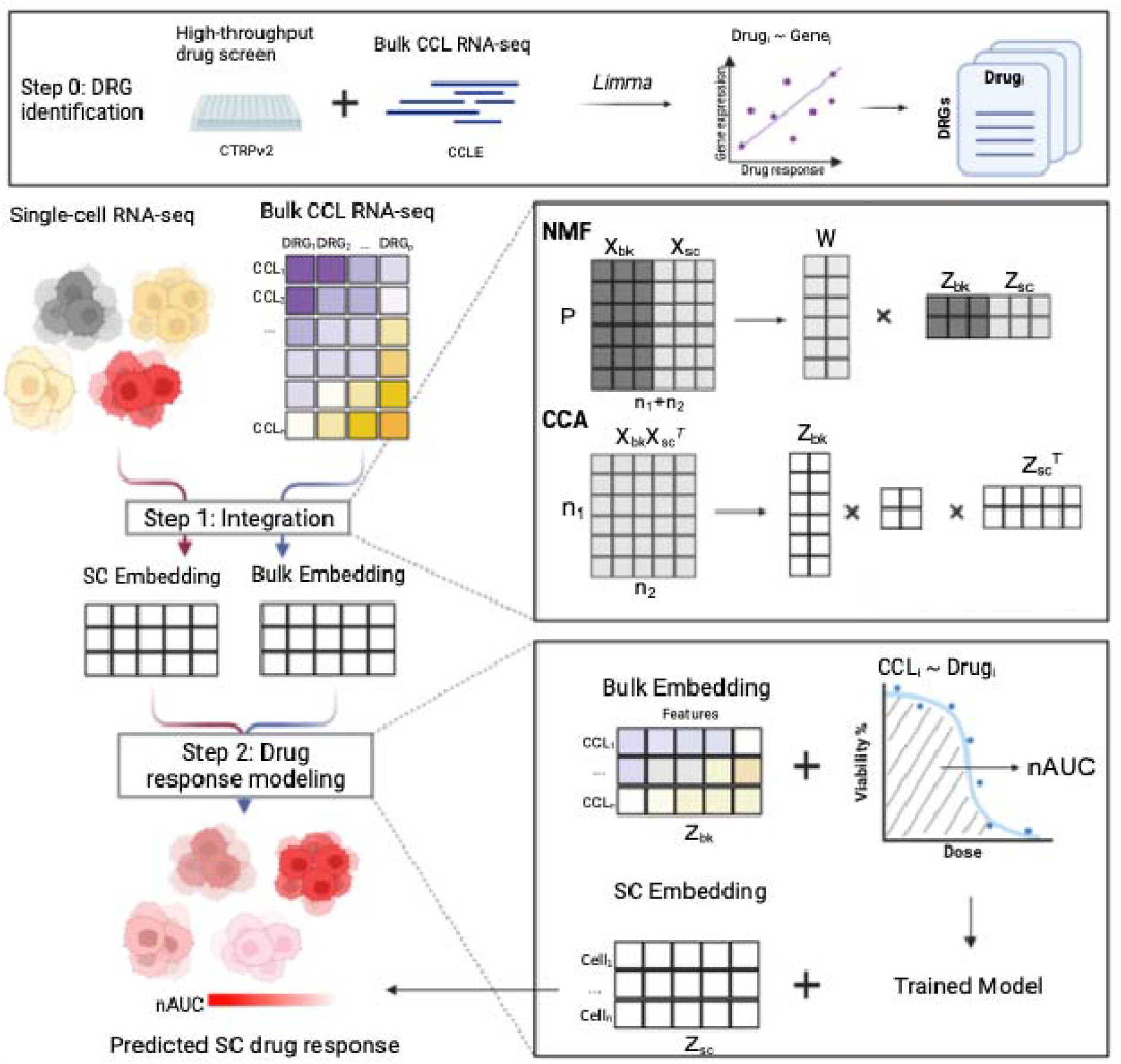
Schematic overview of scIDUC. Drug response relevant genes (DRGs) are generated for use in the scIDUC pipeline (Step 0). Step 1 integrates input CCL bulk RNA-seq data ( ) and scRNA-seq data ( ) to preserve shared expression patterns while reducing noise. The resulting embeddings of bulk RNA-seq ( ) are used with CCL drug response to construct regression models in Step 2. Learned coefficients are applied to scRNA-seq embeddings ( ) to infer cellular drug sensitivity scores. DRG: drug response relevant gene. CCL: cancer cell line. SC: single cell. NMF: nonnegative matrix factorization. CCA: canonical correlation analysis. nAUC: normalized area under the dose response curve.

Given the distinct properties between bulk RNA-seq and scRNA-seq data, the next step is to integrate the CCL bulk RNA-seq dataset and the target scRNA-seq dataset, while preserving shared gene expression patterns and parsing out less relevant noise (<u>Step 1</u>). The rationale of imposing DRGs is to maximize the likelihood of retaining shared transcriptomic patterns that are associated with drug response. Many data integration methods have been proposed to merge multiple scRNA-seq datasets. The scRNA-seq analysis R package Seurat has incorporated canonical correlation analysis (CCA) as one of the core algorithms to combine multiple SC datasets, based on the rationale that CCA will preserve similarities between data sources^30^. Also, non-negative matrix factorization (NMF) has been used for joining scRNA-seq datasets, partially given the interpretation of its inner decomposition factors as “metagenes”^31,32^. Thus, in Step 1, we examined both CCA and NMF algorithms for integrating bulk and SC datasets, which in theory contain less commonalities compared with merging SC datasets only. We also designed experiments to further evaluate performances of CCA and NMF for accurate drug response predictions.

The integration step generates embeddings of the two input RNA-seq datasets and projects them into a low dimensional space. Next in Step 2, we utilized regression-based approaches to model drug response. We trained models using CCL (bulk) embeddings as predictors and measured drug response as the dependent variable. Coefficients of the subspace features were then applied to SC embeddings to generate cellular drug response. This results in predicted nAUC values for all cells in the target scRNA-seq data.

### Selection of parameters and evaluation of pipeline performances

Based on the overall design of scIDUC, one crucial parameter is the number of DRGs, as it directly affects data integration and thus drug response performances. Since the input bulk RNA-seq data mostly remains invariant (CCL expression profile) while the target scRNA-seq varies, we sought to determine this parameter in a data-dependent way, codifying this parameter as the ratio between number of SCs and number of DRGs (or SC-DRG ratio). In addition, the flexible pipeline also allows us to evaluate different means of integration (Step 1) and drug response modeling (Step 2).

Thus, to establish an optimal structure of our pipeline, we applied scIDUC with different parameters or settings to three independent scRNA-seq datasets with known sensitivity status to specific drugs. To be broadly applicable, we chose datasets that represent various diseases and biological origins. In these data, drug resistance was established through chronically exposing model system(s) to drugs of interest, followed by scRNA-seq of both parent drug-sensitive and derived drug-resistant model(s). Specifically, in the Lung-PC9 dataset, Kong et al. chronically exposed PC9 lung cancer cells to Gefitinib^33^ to establish resistance. In the second dataset (Breast-MCF7)^34^, Ben-David et al. developed Bortezomib resistant cells derived from the MCF7 breast cancer cell line; Bell et al. generated cells resistant to BET inhibitors from murine acute myeloid leukemia patient derived xenografts (AML-PDX) models (Supplementary Table S1)^35,36^. We compared predicted cellular drug response from scIDUC with the true sensitive/resistant labels. Specifically, we designed two experiments investigating impacts from (1) means of data integration and drug response modeling as well as from (2) the SC-DRG ratio parameter.

Predicted cellular drug sensitivities from scIDUC were evaluated via calculating two effect sizes between the true resistant and sensitive groups, namely the common-language effect size (Rho statistic) and the Cohen’s D effect size. Rho statistics indicate the probability of a randomly selected cell from the true resistant group having higher predicted nAUC than a randomly selected cell from the true sensitive group (see Methods). A value less than 0.5 indicates contradictory predicted drug response status, a value around 0.5 implies random chance, and a value above 0.5 is ideal. Similarly, Cohen’s D describes differences between predicted nAUCs between the two groups while considering variability (see Methods). A Cohen’s D can generally be interpreted as having a small effect size at 0.2, a medium effect size at 0.5, and a large effect size at 0.8^37^. Higher values of either criterion therefore signify better performances.

#### Selection of integration methods (CCA or NMF) and drug response modeling strategies

We first probed different formulae in both data integration (Step 1) as well as in modeling drug response (Step 2). For integration, given that one crucial parameter for both CCA and NMF is the inner dimension *k* (number of latent factors for NMF and number of canonical correlation vectors, or CCVs, for CCA), we conducted extensive investigations into the robustness of each integration approach with varying *k* values (*k* = {1,2,…50}). Further, for drug response modeling, we incorporated linear regression (Lm) and non-used the first *k* CCVs (for CCA) and the first *k* latent factors (or metagenes, for NMF) respectively and parametric regression models based on a Gaussian kernel (Kernel) to ascertain the optimal pipeline. We examined predicted single-cell drug sensitivities against the truth (Supplementary Figure S2). Two-sample t-tests using predicted nAUCs were performed between the true resistant cells and sensitive cells, based on which a positive t-statistic indicates correct predicted directions. For Lung-PC9, both methods were able to generate correct sensitivity trends towards gefitinib, while CCA based integration shows superiority in terms of p-values and t-statistics. For Breast-MCF7 and AML-PDX, CCA consistently predicted correct cellular drug response status, whereas volatile test statistics were observed with NMF, especially NMF with downstream linear models for drug response prediction. Although NMF coupled with kernelized regression resulted in more stable results than NMF and linear models, CCA continued to show superior results regardless of regression models (Supplementary Figure S2). Taken together, CCA integration showed superiority over NMF considering both accuracy and robustness. We also found that nonparametric modeling of drug response seemed to work better for NMF compared with linear models. When coupled with CCA, both regression models gave comparable results. Additionally, while the selection of the optimal *k* in either CCA or NMF plays a central role in algorithm performance^30,38^, given that the computational goal is to model drug response, we implemented feature selection on post-integration embeddings to include subspace features that correlate with drug response (see Methods). Through this, scIDUC was able to quickly select only a few meaningful features for model training and prediction without searching for optimal inner dimensions in an unsupervised fashion.

#### Identification of optimal SC-DRG ratios

Given results from (1), we further tested varying SC-DRG ratios when using (a) no integration, (b) CCA+Lm, and (c) NMF+Kernel. An SC-DRG ratio range spanning from 10 to 0.2 was examined. For each SC-DRG, in Step 2, a subset of 95% of available CCLs were randomly selected as a bootstrap sample to train prediction models. This process is repeated 50 times, which allowed us to test scIDUC’s stability and robustness. Evaluation results were reported at each SC-DRG ratio (Figure 2). When integration was performed, median Cohen’s D surpassed 0.8 and median Rho statistics surpassed 0.5 for the majority of ratios tested, implying that scIDUC can in general recapitulate known cellular response to drugs across three scRNA-seq datasets regardless of SC-DRG ratios. No clear trend was observed from results without integration. Predicted nAUCs with integration also showed higher robustness, indicated by less variability in the results. While with both integration algorithms the prediction accuracy tends to peak when SC-DRG ratios fell between 1 and 0.2, NMF showed higher variability compared to CCA and had poorer performances outside this ideal SC-DRG ratio range (Figure 2 and Supplementary Table S2). We observed that a SC-DRG ratio between 1 to 0.2 in general gave good results, supported by stable Rho statistics close to 1 and Cohen’s D larger than 1 across all three scRNA-seq data.

**Figure 2.**
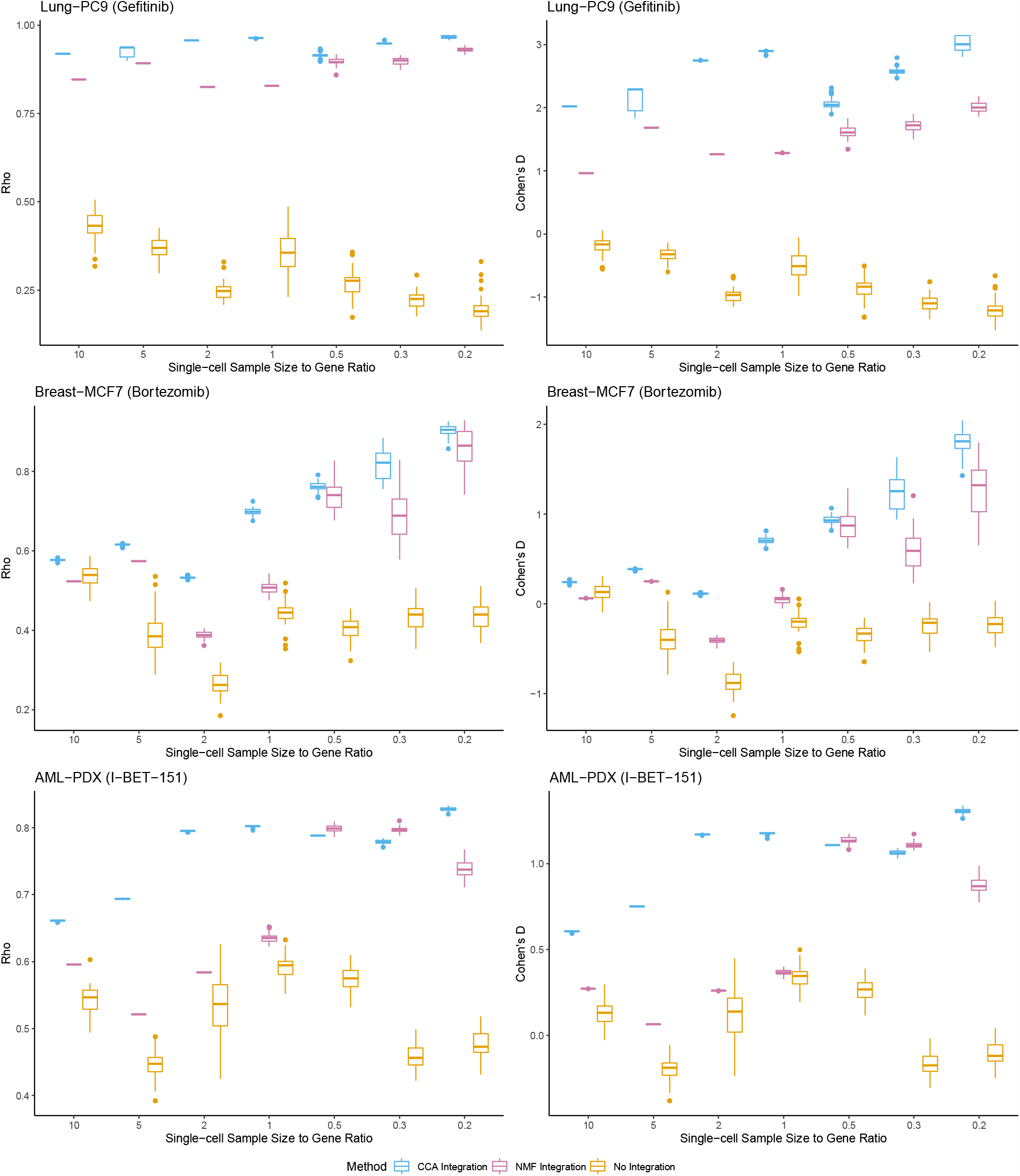
scIDUC recapitulates cellular drug sensitivity status in scRNA-seq data. We applied scIDUC with different data integration settings using varying SC-DRG ratios. Distribution of common-language effect Rho (left column) and Cohen’ D (right column) comparing predicted cell response between the true resistant vs. sensitive cell groups across three scRNA-seq datasets are shown. A larger value indicates a higher separation between the groups. Top row: Lung-PC9; mid row: Breast-MCF7; bottom row: AML-PDX. Basic information of these dataset can be found in Supplementary Table S1. Summarized mean and standard deviation of the effect sizes are shown in Supplementary Table S2.

Taking results from (1) and (2) together, we prioritize CCA as scIDUC’s core integration method. We employ linear regression to model drug response for its simplicity and interpretability. A SC-DRG ratio between 0.2 and 1 was recommended for obtaining optimal predictions. However, the final build of scIDUC package does allow users to explore NMF and kernel regression as alternative settings.

### scIDUC outperforms other methods in scRNA-seq data from various sources

Next, scIDUC was benchmarked against other methods, namely Beyondcell^22^ and CaDRReS-Sc^21^, which also aim to predict cellular drug response (Supplementary Information 1). Apart from the three datasets used to evaluate the scIDUC pipeline, we included three additional scRNA-seq data, representing various biology backgrounds as benchmarking datasets (Supplementary Table S1). In CRPC-CCLs, Schnepp et al. exposed castration-resistant prostate cancer (CRPC) cell lines PC3 and DU145 to incremental doses of docetaxel to acquire resistance^39^. The PDAC-CFPAC1 dataset describes ductal pancreatic adenocarcinoma CFPAC1 cells whose drug response was profoundly altered by tumor microenvironment (TME)^40^. When growing in complete classical organoid media, CFPAC1 cells lost basal properties and became responsive to several treatments including SN-38 and paclitaxel. In RMS-oPDX, Patel et al. discovered a mesoderm-like cell colony using scRNA-seq on orthotopic patient-derived xenografts (oPDX) from pediatric patients with rhabdomyosarcoma (RMS). These cells were shown to be highly resistant to the chemotherapy irinotecan (whose active metabolite is SN-38) compared to myoblast cells but sensitive to EGFR inhibitors^41^. These data allowed us to investigate performances of scIDUC and competing methods in diverse biology models and indications.

We benchmarked prediction performances of scIDUC, Beyondcell^22^, and CaDRReS-Sc^21^ using the same evaluation criteria, namely Cohen’s D and Rho statistics. Bootstrapping was implemented for all three methods (see Methods). For scIDUC, since a [0.2,1] SC-DRG range generally produced good results based on our experiment results from Figure 2, we used two SC-DRG ratios (0.2 and 0.9) within this range to demonstrate the minimal parameter tuning needs for scIDUC, while avoiding biased results. Across all datasets, our method achieves the highest effect sizes across all three benchmarking datasets under both SC-DRG specifications, with median Rho-statistics above 0.8 and median Cohen’s D above 1 (Figure 3). Beyondcell was able to separate the true resistant and sensitive cells and showed high consistency, however its results were less accurate (median Rho-statistics around 0.7 and median Cohen’s D less than 1) compared with scIDUC. CaDRReS-Sc did not generate meaningful predictions to recollect cellular drug response (median Rho-statistics around 0.5 and median Cohen’s D close to 0). Additionally, across datasets Lung-PC9, Breast-MCF7, and AML-PDX, we observed similar results: scIDUC in general had the highest Rho statistics and Cohen’s D; Beyondcell provided meaningful predictions though less accurate. Interestingly, CaDRReS-Sc showed good performance with the Breast-MCF7 dataset, though failed to recapitulate true cellular drug response status in the other two. Notably, for AML-PDX, neither Beyondcell nor CaDRReS-Sc was able to recollect true cell drug response status (Supplementary Figure S3). The suboptimal performances of the two other methods were not unanticipated given their methodological designs. For Beyondcell, though applying bulk-learned signatures to SC data could bypass integrating bulk and SC datasets and to some extent reflect cellular response to drugs, these signatures are swayed by quality of scRNA-seq data. Signature score calculating can be subordinate to random factors such as drop-outs and low expression values, resulting in less ideal predictions. CaDRReS-Sc centers its modeling strategies around IC50 instead of AUC that has been demonstrated to be more reliable for response prediction^42^. Here, our benchmarking results could suggest that modeling strategies used by CaDRReS-Sc lack adaptivity to account for AUCs as drug response. Furthermore, a fixed set of drug response essential genes were used by CaDRReS-Sc for all drugs^43^. Since different drugs might correlate with different genes, an invariant gene set might be insufficient to capture broad drug-gene relationships. In comparison, scIDUC and Beyondcell are both sensitive to drug-specific genes and showed superior results.

**Figure 3.**
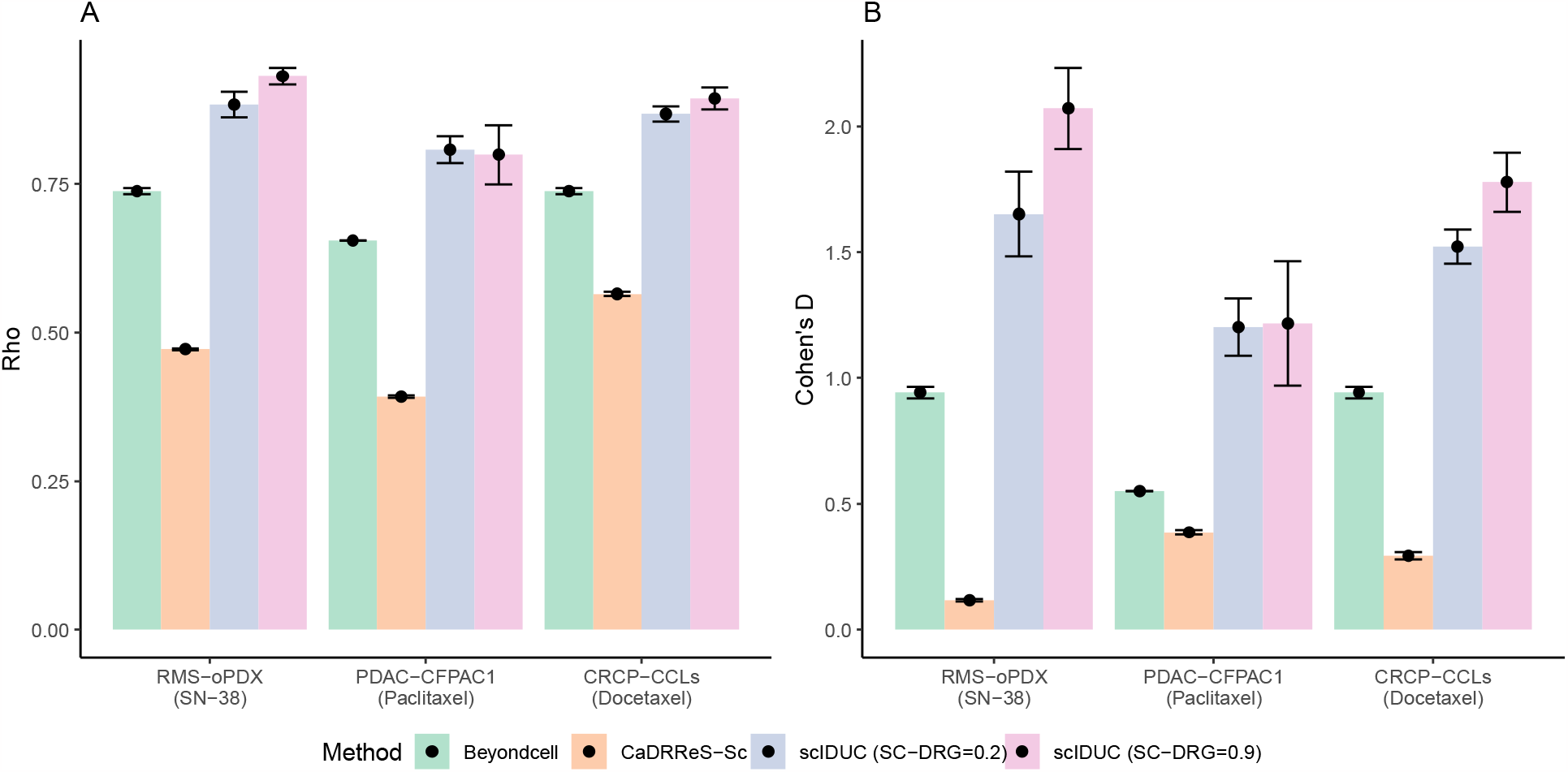
scIDUC outperforms other methods across three scRNA-seq datasets. Mean and standard deviation of Rho statistics and Cohen’s D are shown for all three methods across the benchmarking datasets. For each method, 50 bootstrap samples were generated (see Methods). In all three datasets, scIDUC shows higher Common-language effect Rho (A) and Cohen’ D (B) comparing predicted cell response between the true resistant vs. sensitive cell groups than that of other methods (CaDRReS-Sc or Beyondcell). Summarized numeric results are shown in Supplementary Table S3.

### scIDUC enables clinically meaningful drug discovery

We next showcase the utility of our scIDUC framework as a trailblazer in aiding hypothesis development for clinically meaningful drug discovery. We conducted prospective analyses for three different scenarios utilizing the RMS-oPDX, PDAC-CFPAC1, and CRPC-CCLs datasets, representing diverse biology models including patient-derived xenografts (PDX), tumor microenvironments (TME), and acquired drug resistance *in vitro*. In RMS-oPDX, we applied scIDUC to screen efficacious drugs targeting the SOC (SN-38) resistant mesoderm-like cells. Our nominated drugs showed high consistency with findings from the original study. In PDAC-CFPAC1, scIDUC was used to predict sensitivities of CFPAC1 cells grown in different TMEs to various drugs. The resulting differential efficacy profile between the two TMEs were highly comparable to drug panel results reported by Raghavan et al. Finally, in CRPC-CCLs, we utilized scIDUC to screen drugs showing efficacies in docetaxel-resistant CRPC cells and successfully validated our nomination through in vitro experiments.

#### Nominating drugs targeting mesoderm cells in RMS patients

In RMS-oPDX, Patel et al. delineated that mesoderm-like cells in pediatric RMS tumors bore profound resistance to SOC chemotherapy irinotecan (SN-38)^41^. Therefore, targeting therapy-resistant mesoderm cells constitutes a cornerstone for curbing the current high rates of disease recurrence. To this end, we applied scIDUC to RMS-oPDX, aiming to discover drugs showing high efficacy in mesoderm cells. To increase likelihood of finding actionable therapeutics, we expanded the pool of candidate drugs by predicting cellular sensitivities to various compounds from not only the CTRPv2 but in addition the Genomics of Drug Sensitivity in Cancer 2 (GDSC2)^44^ databases. Two-sample testing was performed between mesoderm (SOC resistant) cells and myoblast (SOC sensitive) cells, through which drugs displaying lower predicted nAUC (suggesting higher sensitivity) in mesoderm cells were included as candidates. To ensure prediction robustness, this pipeline was applied independently to oPDX from each patient (11 in total) in the dataset, and frequency of each nominated drug was summarized over all patients. We considered drugs with a frequency higher than 50%, or nominated from at least 6 out of 11 patients independently as robust candidates. We identified drugs with diverse mechanisms and ranked their target pathways by the number of drugs belonging to the same class (Figure 4A). In both CTRPv2 and GDSC2, epidermal growth factor receptor (EGFR) is among the most frequent targets (Figure 4A). Furthermore, 16 drugs were simultaneously proposed to be efficacious against mesoderm cells by comparing prediction results from both databases. The top five target pathways of the 16 identified compounds were presented in Figure 4B, in which EGFR is ranked the first, targeted by three drugs (afatinib, gefitinib, and erlotinib). Our drug nomination is strongly supported by the original study, where EGFR was validated as an actionable drug target for chemo-resistant mesoderm cells. The application of scIDUC recapitulates such a finding through independent, data-driven analysis. In addition, scIDUC has proposed other drugs and target pathways as potential strategies for inhibiting mesoderm cells in RMS patients, such as MEK inhibitors (trametinib and selumetinib). MEK inhibitors were previously shown to induce tumor differentiation in RMS and potently inhibit RMS in both cell line models and xenograft models^45,46^. These findings warranted the further evaluation of these newly nominated drugs in SOC resistant RMS patients.

**Figure 4.**
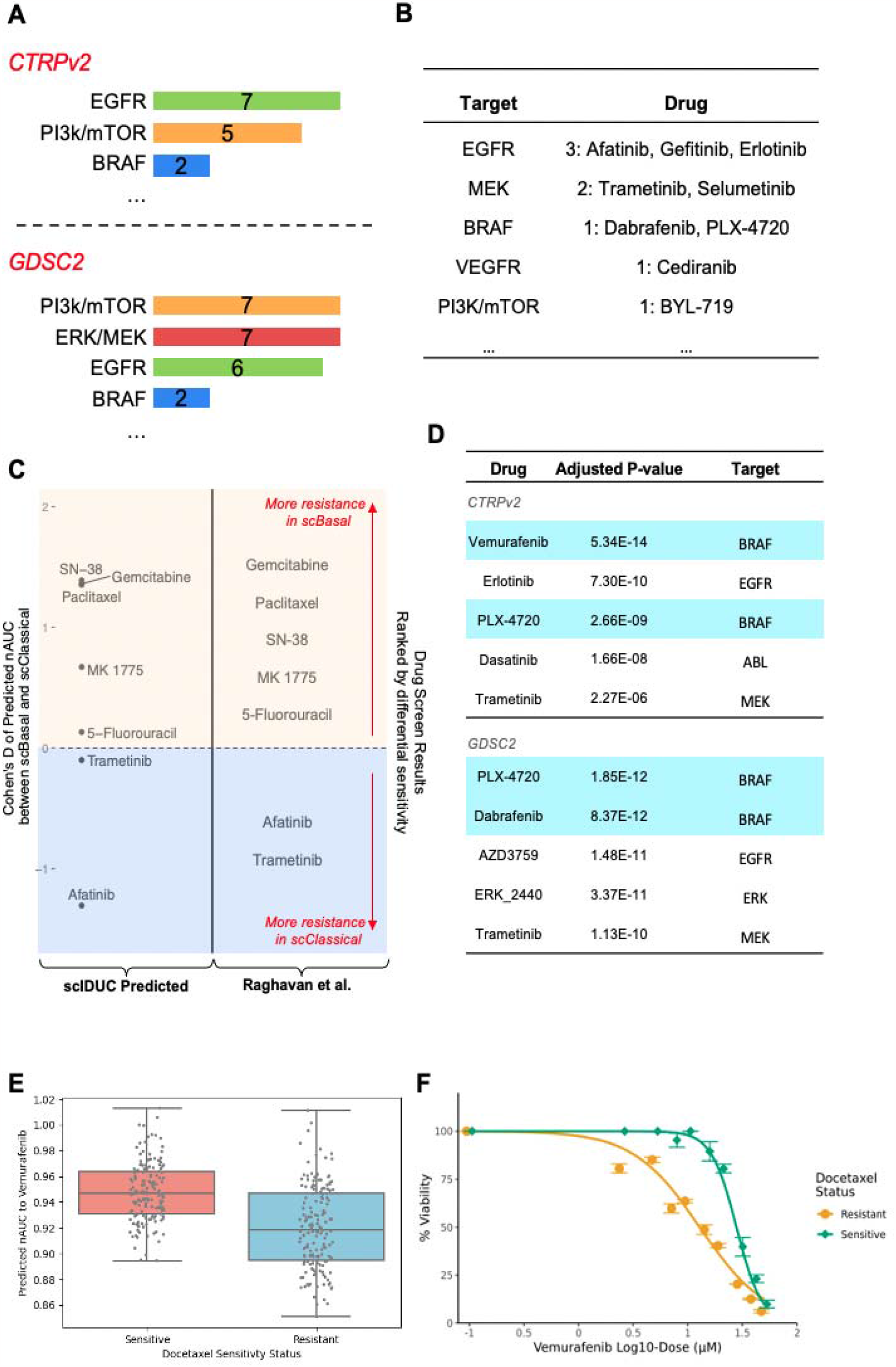
scIDUC facilitates drug discovery in various models by identifying cell-type-specific drug candidates. **A**. Top drug targets predicted by scIDUC for SOC-resistant mesoderm-like cells in RMS-oPDX. **B**. Efficacious drugs for mesoderm-like cells in RMS-oPDX concurrently identified from the CTRPv2 and the GDSC2 and their targets. **C**. scIDUC recapitulates differential efficacies shaped by tumor microenvironments (TME) in the PDAC model CFPAC1 cells. **D**. Top candidate drugs predicted to be efficacious against docetaxel resistant cells from each high-throughput drug screen **E**. BRAF inhibitor vemurafenib is predicted to be effective against docetaxel-resistant DU145 cells (adjusted p<0.0001). For each cell group, the box shows the median, the first, and the third quartiles of predicted nAUCs. **F**. Docetaxel-resistant DU145 cells show higher sensitivity to vemurafenib compared to their docetaxel-sensitive counterparts in vitro (two-way ANOVA p<0.0001). At each concentration, mean percent viability ± standard deviation is plotted. SOC: Standard-of-Care. CTRPv2: Cancer Therapeutic Response Portal Version 2. GDSC: Genomics of Drug Sensitivity in Cancer. PDAC: Pancreatic Ductal Adenocarcinoma.

#### Depicting therapeutic susceptibility in PDAC cells affected by different TMEs

Raghavan et al. showed that TME dramatically altered sensitivities to various therapeutics in the PDAC cell line model CFPAC1 as well as in PDAC organoids^40^. For example, as previously recapitulated by scIDUC, CFPAC1 cells grown in classical complete organoid media (scClassical) were re-sensitized to SN-38 and Paclitaxel compared to their counterparts grown in the basal cell line media (scBasal). An additional measured drug panel screen was performed by the authors to obtain a broader differential drug response profile between scBasal and scClassical states induced by different TMEs (Supplementary Figure S4A, or originally Figure 6G from Raghavan et al.). To study predictability of scIDUC in this situation, we predicted sensitivities of scBasal and scClassical cells to the same treatments tested in the original drug panel. Based on the drug panel, treatments with demonstrated differential efficacy (i.e., treatments whose mean differential efficacy deviated from zero and confidence interval did not contain zero) were kept, among which seven drugs were found in the CTRPv2 database and used by scIDUC to generate cellular response predictions. Two-sample t-tests were conducted to examine differences in predicted nAUCs between scBasal and scClassical cells (Supplementary Figure S4B). Cohen’s D was calculated to demonstrate differences in predicted drug response between scBasal and scClassical cells; a positive Cohen’s D indicates more resistance in scBasal cells, whereas a negative value implies the contrary. We plotted Cohen’s D of each treatment by its actual value to illustrate the extent of predicted differential efficacy between the two cell states (Figure 4C, left). Corresponding drug panel results from Raghavan et al. were shown in their original ranking for comparison (Figure 4C, right). Our results achieved high consistency with drug panel data by Raghavan et al. Gemcitabine, SN-38, and paclitaxel had the highest Cohen’s D, indicating profound discrepancies between resistant scBasal cells and sensitive scClassical cells. MK 1775 and 5-Fluorouracil each showed modest and slight differences, whereas trametinib and afatinib were predicted to be more resistant in scClassical cells. Our results were strongly supported by the original experimentally measured drug response in different TMEs, which highlights the capability of scIDUC at capturing TME-shaped drug response at the single-cell level. Furthermore, scIDUC can be applied to hundreds of drugs screened in CCL drug screenings to decipher potential differential drug response in different TMEs, which is extremely expensive if not impossible to carry out at this scale experimentally.

#### Discovering and validating drugs for docetaxel-resistance in CRPC

Though docetaxel was approved for CRPC, resistance among patients is prevalent^39^, highlighting a need to identify new therapeutics targeting the non-responsive cells. We utilized the CRPC-CCLs dataset where docetaxel sensitive and resistant cells have been experimentally defined and supported by our scIDUC prediction. Here we employed scIDUC again to predict cellular sensitivities to hundreds of other treatments in the CTRPv2 and the GDSC2 databases and prospectively identified drugs showing predicted efficacy in docetaxel resistant cells. We conducted two-sample t-tests comparing docetaxel resistant and sensitive cell groups, through which we selected drugs showing higher effects (lower nAUCs) in the resistant group using an adjusted p-value threshold of less than 0.05. Resulting drugs from each database were ranked based on their adjusted p-values from the lowest to the highest. We selected the top five drugs with the smallest adjusted p-values from each data source and listed their molecular targets (Figure 4D). A number of BRAF inhibitors were identified from the CTRPv2 (vemurafenib and PLX-4720) and the GDSC2 (PLX-4720 and dabrafenib). In CTRPv2, vemurafenib showed the highest differential efficacy between the two CRPC cell groups, with docetaxel resistant cells having significantly higher predicted sensitivity than docetaxel resistant cells (Figure 4E, adjusted p-value=5.34 x 10^-14^). To experimentally validate this prediction, we chose to test vemurafenib using a previously developed *in vitro* cell line model system containing docetaxel sensitive and resistant DU145 cells. We first exposed these established cells to docetaxel to ensure presence of differential response to the drug (Supplementary Figure S5). To evaluate our candidate drug, we treated cells with increasing concentrations of vemurafenib and generated dose-response curves for both cell groups (Figure 4F). Vemurafenib showed significantly higher inhibitory activity among docetaxel resistant DU145 cells with a half maximal inhibitory concentration (IC50) of 12.9 μM/L compared to its IC50 of 27.9 μM/L in sensitive cells (Figure 4G, two-way ANOVA p<0.0001). Taken together, our *in vitro* experiment results matched our computational predictions, further supporting the reliability of prospective results generated by scIDUC. Overall, through applying scIDUC in three different scenarios fulfilling varying research needs, we demonstrate its ability to enable drug development targeting heterogeneous tumors. Further examination and experimental validation of our prediction results underscore the potential clinical impact of identified drugs. The diverse biological backgrounds in these cases, including PDX, TMEs, and *in vitro* cell models, support broad adaptations of scIDUC to enable clinically meaningful drug discovery.

## Discussion

Assessment of gene expression at the SC level offers detailed mappings of cell compositions and substantially advances understanding of complex diseases such as cancer that involves heterogeneous cell types. Origins of therapy resistance and disease recurrence have been linked to such heterogeneity in many malignancies, often attributed to existence of insusceptible cells thriving under selective pressures^1–3^. Accordingly, cell-type-aware drug discovery using scRNA-seq data has demonstrated potentials to curb resistance and improve treatment outcomes^47–49^. Though computational drug discovery tools have been proposed for this goal amid increasing public access to scRNA-seq datasets, most pipelines still focus on target identification and validation, a procedure often involves generation of scRNA-seq data, or inference of cellular changes under therapeutic perturbations (instead of cellular response to a potential treatment given its expression profile)^10,50^. These methods so far have not been demonstrated to be able to utilize existing scRNA-seq data to conduct virtual drug screens, facilitate hypothesis formulation, and propose drug candidates addressing tumor heterogeneity. To fill this research need, we have developed scIDUC, which integrates the pan-cancer CCL bulk RNA-seq and scRNA-seq data and infers cellular response to various drugs. By coercing both datasets to have the same DRGs, the CCA-based integration preserves similarities between bulk and scRNA-seq data in a drug response relevant context, allowing accurate predictions to be made at the single cell level without the need for scRNA-seq drug screen training data, which is difficult to acquire. More importantly, through prospective analysis and validation in three distinct scenarios, we demonstrated the versatility of scIDUC for quickly generating cell-type-specific predictions. The resulting predictions showed high concordance with previous findings and experiment results, further bolstering the utility of scIDUC for providing therapeutic opportunities with clinical impact.

To configure the scIDUC pipeline for optimal results, we evaluated its performances with varying factors including SC-DRG ratios, integration methods, and drug response models against known cell drug response status in independent scRNA-seq datasets. Our results spotlighted an SC-DRG ratio range between 0.2 to 1, data integration via CCA, and linear regression models for accurate predictions. As we constructed and evaluated scIDUC using CTRPv2 as source for drug-gene information extraction, we recommend the use of CTRPv2 as default drug screen database for drug discovery. Nonetheless, our framework also includes other high-throughput screens such as the GDSC2 and the Genentech Cell Line Screen Initiative (gCSI) if needed by the user. Since scRNA-seq data can consist of parental and resistant cell populations receiving different treatment, we gathered additional evidence showing that scIDUC predictions were not confounded by potential batch effects. First, we compared predicted sensitivities to a variety of drugs between true resistant and true sensitive cell groups in AML-PDX (Supplementary Figure S6). Considering the established resistance against I-BET-151, predicted cellular response toward another BET inhibitor, I-BET-762, also showed significant differential sensitivity. However, predicted response to other drugs such as histone deacetylase inhibitors (entinostat and vorinostat), NAMPT inhibitor (daporinad), and EGFR/HER2 inhibitor (lapatinib) showed minimal to no separation between the two groups. Moreover, in the RMS-oPDX dataset, cells from an oPDX sample underwent sequencing altogether and therefore precluded batch effects. In this scenario, scIDUC not only recapitulated resistance to the SOC therapy SN-38 but provided drug nomination highly in line with the original findings. Collectively, our results show that scIDUC predictions are not hindered by potential existence of batch effects.

To date, a few other methods have been proposed to computationally predict single cell drug response using drug screen data on CCLs, such as Beyondcell, CaDRRes-sc, SCAD, and scDEAL^21–23,26^. Each of these methods employ a unique transfer learning-like approach, utilizing relationships between CCL expression data and drug response to predict single cell level drug sensitivity. However, there are several differences in the actual methodology among these methods, which is reviewed in Supplementary Information 1. We compared scIDUC with CaDRRes-sc and Beyondcell and demonstrated its superiority in prediction accuracy. Across the three independent benchmarking scRNA-seq datasets (RMS-oPDX, PDAC-CFPAC1, and CRPC-CCLs) with known cellular drug response labels (gold standards), cellular drug sensitivities predicted by scIDUC not only reiterated true drug response but also had highest precision (evidenced by highest rho statistics and Cohen’s D). In comparison, results by Beyondcell echoed true cellular drug sensitivities but lacked accuracy, whereas CaDRRes-sc in general failed to provide meaningful predictions (Figure 3). To be comprehensive, we have conducted the same analysis using the first three datasets (Lung-PC9, Breast-MCF7, and AML-PDX) and observed similar trends for each. Since we fine-tuned scIDUC parameters using the first three datasets, these results were given in Supplementary Figure S3 and used only as additional supporting evidence. Notably, CaDRRes-sc utilizes an invariant set of essential genes generated from CRISPR screens in its data integration process^43^, while both scIDUC and Beyondcell derive drug-specific marker genes. Our benchmarking results support the rationale of using genes whose expression correlates with measured drug response. Since CRISPR screens detect genes altered by therapeutic perturbations, they may not reflect drug-gene relationships at the baseline level. On the other hand, using signatures alone is susceptible to varying quality of scRNA-seq data which are known to have low detection rates. Meanwhile, calculated scores based on these signatures reflect only relative sensitivities within a scRNA-seq data and lack pharmacological meanings. Taken together, scIDUC achieves desirable results through incorporating both drug-specific features and bulk-SC integration. In addition, SCAD and scDEAL both employ DL methodologies to perform bulk-SC integration and drug response prediction. Both methods embody binarized labels as drug response and train corresponding models as classification problems. Given that drug response across CCLs is better described by spectrum values such as AUCs^51^, it is unclear if binary cellular drug labels can capture varying degrees of sensitivity in a heterogeneous tumor. Furthermore, insufficient evidence was given by competing methods to demonstrate how resulting SC drug response will benefit hypothesis generation and drug discovery. It is also noteworthy that compared with other methods, scIDUC requires minimal parameter tuning, enabling adaptations to a broad user base for various oncology therapy research topics.

Successful characterization of drug response profiles at the SC level plays a fundamental role in advancing precision medicine in cancers^52,53^. Learned cellular sensitivity to various drugs can greatly benefit studies tackling topics such as heterogeneity and cancer drug resistance by providing cell type specific vulnerability information. Aided by the robust prediction results from scIDUC, we spearheaded hypothesis generation and drug candidate identification in three distinct scenarios. For RMS-oPDX, we applied scIDUC independently in 11 oPDX samples to identify drugs showing efficacy against the SOC therapy resistant mesoderm-like cells. In the majority of the samples EGFR inhibitors were nominated as one of the potential drug classes, reiterating original findings from Patel et al^41^. Moreover, we also discovered a number of other drug classes as potential targets. For example, MEK inhibitors showed high occurrences combating mesoderm-like cells. Previous studies have established evidence that MEK inhibitors effectively inhibit RMS both *in vitro* and *in vivo*^45,46^. Moreover, inhibition of the MAPK/ERK pathway by MEK inhibitors have been shown to downregulate mesodermal genes in embryonic stem cells^54^. Given these findings, further evaluation of MEK inhibitors in RMS patients with disease recurrences is warranted. In the second scenario, we showcased that scIDUC was able to capture TME-shaped differential drug response in CFPAC1 cells, a PDAC cell line model. Microenvironmental niche factors have been shown to drive aggressive PDAC progression from a therapy-responsive “classical” state to a less differentiated “basal” state^40,55^. Thus, characterizing PDAC cellular drug response in different TME-driven states is a crucial first step to develop treatments to curb the current high mortality rate (five-year survival ∼9%). Differential cellular drug efficacies between scBasal and scClassical cells resulted from scIDUC accurately recapitulate drug panel testing results reported by Raghavan et al. (Figure 4C). In addition, Shinkawa et al. also reported similar findings where basal-like, poorly progressed PDAC organoids showed higher resistance to gemcitabine compared to classical-like organoids^55^. These discoveries substantially support usage of scIDUC to streamline drug discovery under different TMEs without the need to conduct large scale drug screens. Finally, we utilized scIDUC again to screen for alternative therapeutics against docetaxel resistance in CRPC CCLs including DU145 and PC3. Our top candidate, namely the BRAF inhibitor vemurafenib, showed consistent higher efficacy in docetaxel resistant DU145 cells than sensitive ones when evaluated *in vitro* (Figure 4F). A previous trial of vemurafenib has reported that an average maximum serum concentration (Cmax) of 61.4 μg/mL, or equivalent to 125.3 μM/L, was well tolerated among patients^56^. Our in vitro experiments exposing vemurafenib in docetaxel resistant cells estimated an IC50 of 12.9 μM/L, which sits well below the safety dose, highlighting its clinical potential to be used in combination with docetaxel to control CRPC progression. Furthermore, two EGFR inhibitors have been proposed to combat docetaxel resistance by our computational pipeline (Figure 4D), consistent with previous studies suggesting that EGFR inhibitors mediate docetaxel resistance in CRPC^57,58^. To sum up, our scIDUC powered computational pipelines were able to quickly propose drug candidates with clinical impact. Our method was able to pinpoint a selective collection of actionable drugs that are ready to be evaluated for different purposes. Serving as an alternative to traditional drug screens and target identification, scIDUC was able to provide rationalized and streamlined drug nomination for therapeutic development. When combined with experimental testing, it can speed up development of efficacious treatments.

In addition, knowledge of intratumoral therapy vulnerability has been explored to inform formulation of drug combinations that target multiple cell groups to help eliminate heterogeneous tumors^20,59^. Given the vast number of potential combination therapies, computational frameworks have been proposed to conduct virtual systematic screens for specific indications^60,61^. To this end, cellular drug response scores are key components for modeling combination efficacies. For example, nAUC might be perceived as the probability of an organism surviving a certain drug treatment; under such an assumption, cellular nAUCs can be used to infer potential drug synergy under various statistical models^62^. Cellular drug response scores from scIDUC provide key variables for drug combination modeling strategies; our future work will incorporate scIDUC and computational drug combination discovery pipelines to establish a virtual screen platform for various complex cancers.

In sum, we showcase a computational method to depict SC vulnerability to various drugs, establishing a foundation for cell-type-aware drug discovery combating the prevailing issue of treatment failure due to tumor heterogeneity. Our case studies not only provide therapeutic options for various diseases but also substantiate the necessity of our proposed method in aiding efficient development of clinical meaningful treatments.

## Methods

### Data acquisition

The pan-cancer cell line (CCL) transcriptomic data (bulk RNA-seq) was downloaded from the Cancer Dependency Map (DepMap, https://depmap.org/portal/)^63^ and originally from the Cancer Cell Line Encyclopedia (CCLE)^16^. CCLE expression data were downloaded in raw count and log2(TMP+1) formats. CCL drug response data was downloaded from DepMap, originally from the Cancer Therapeutics Response Portal (CTRPv2) generated at the Broad Institute^15,63,64^. We utilized raw drug screen data to refit dose-response curves and retain robust drug sensitivity profiles^65^. For each drug-CCL pair, area-under-the-dose-response-curve (AUC) was divided by its tested dose range to generate normalized AUC (nAUC), which was then used as the drug response in scIDUC; nAUC is continuous and ranges between 0 to 1 with 0 implying complete cell kill and 1 implying no cell kill.

The single-cell RNA-seq datasets used in this study were downloaded from various sources, depending on the availability of original data provided by the authors. A detailed description of each data source and its properties can be found in Supplementary Table S1.

### Preprocessing

CCL names in downloaded CCLE transcriptome and CTRPv2 drug response data were harmonized to Cellosaurus accession numbers, which make use of the prefix “CVCL”^14,66^. Given that a drug was only screened in a subset of CCLs, we calculated percentages of missing values for each drug and excluded those screened in less than 40 percent of all CCLs in the database. This results in a total of 493 treatments (and 887 CCLs) in the CTRPv2 dataset.

For scRNA-seq data with raw counts, each dataset was pre-processed using the Scanpy Python module^67^. A threshold was imposed on all datasets to filter for cells with at least 200 genes detected and genes detected in at least 3 cells. Each cell was then normalized to have the same total counts of 1 million (counts per million, CPM) and log-transformed with a pseudo-count of 1, i.e., log2(CPM+1).

### Drug response relevant genes generation

We used the R package limma to detect drug response relevant genes (DRGs) given the continuous nature of nAUC. As recommended by the package, for each drug, CCLE raw expression counts were used to construct linear models. Resulting genes were ranked by B-statistics which indicates probabilities of differentially expressed from most significantly associated with drug response to least.

### Integration of bulk and sc data

We implemented two different approaches to integrate bulk and SC RNA-seq data, namely canonical correlation analysis (CCA) and non-negative matrix factorization (NMF). Let 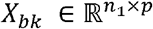 be the CCL bulk RNA-seq matrix with *n*_1_samples and *p* genes; let 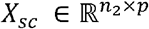 be a scRNA-seq matrix with *n*_2_ cells and genes. The bulk- and SC-matrices have the same DRGs.

#### Integration via CCA

We expanded the CCA integration pipeline proposed in the Seurat package^30^. Briefly, we conducted singular value decomposition (SVD) on the matrix derived based on the multiplication of 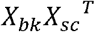. 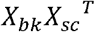 captures similarities between CCLs from bulk RNA-seq and cells from scRNA-seq based on the shared DRGs. Therefore, resulting singular vectors through SVD, i.e.,

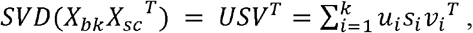

can be viewed as canonical correlation vectors (CCVs). The left singular vectors 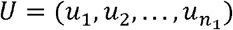 correspond to CCVs for bulk data; the right singular vectors 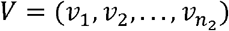 correspond to CCVs for SC data.

Accordingly, 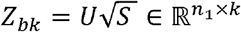 and 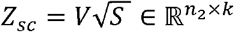 provides embeddings of *X*_*bk*_ and *X*_*sc*_ to a subscape where similarities between bulk and single-cell data are preserved. A visualization of this process is provided in Supplementary Figure S1.

#### Integration via NMF

Since several studies have reported NMF-based methods to capture common gene expression patterns and adjust for discrepancies between batches^31,32^, we included NMF as an alternative means to integrate bulk and single-cell data. Let 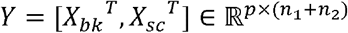 be a concatenated matrix containing both data sources, then

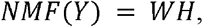

where 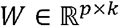 is a common factor matrix whose columns can be viewed as metagenes; 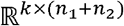 describes metagene expression profiles for bulk samples and single cells. Within 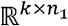 and 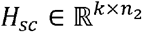 are metagene expression matrices for bulk data and SC data, respectively.

### Drug-response relevant feature extraction

We performed ad hoc feature extraction to select a pharmacogenomic subspace (if CCA) or pharmacogenomic metagenes (if NMF) for accurately inferring single-cell drug response without the need to determine the inner dimensionality within the matrix decomposition tasks. For embeddings resulted from CCA integration, we correlate each dimension (feature) in *Z*_bk_ (bulk embeddings) with measured drug response. Resulting p-values are adjusted via the Benjamini–Hochberg procedure to control false discovery rates (FDRs)^68^. Dimensions that have FDRs less than a threshold are retained. In other words, such pharmacogenomic subspace comprises dimensions *r* = { *r*_1_, *r*_2_,…, *r*_*j*_}⊂ { 1, 2,…, *k*} where *FDR*(*r*_*j*_) < δ and δ ∈ {0.05, 0.1} by default. Training data is then defined as 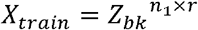, and predictions of cellular drug response will be made on 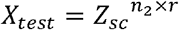.

For NMF, we select metagenes by correlating each metagene in *H*_bk_ with measured drug response. Metagenes that have FDRs less than a threshold π are kept. We retain a set of metagenes *m* = { *m*_1_, *m*_2_,…, *m*_*l*_}⊂ { 1, 2,…, *k*}, where *FDR*(*m*_*l*_) < π. Training data is then defined as 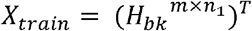, and predictions of cellular drug response will be made on 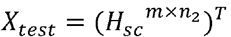.

### Prediction

To predict single-cell drug response, we formulate a linear regression model using *X*_*train*_ as predictors and measured drug response as the dependent variable. Learned coefficients are then applied to *X*_*text*_ to generate nAUCs for cells. We also included an alternative non-parametric regression model based upon the radial basis function (RBF) kernel.

### Evaluation metrics

To evaluate the performances of algorithms, we compared predicted cellular nAUCs against true drug sensitivity status (resistant or sensitive) via two-sample t-tests. To better illustrate, we incorporated two additional metrics showing the effect sizes of predicted drug response differences between resistant and sensitive cell groups.

Let the predicted nAUCs of resistant cells be *L*_*r*_ = (*l*_*r*1_, *l*_*r*2_,…, *l*_*rp*_) and that of sensitive cells be *L*_*s*_ = (*l*_*s*1_, *l*_*s*2_,…, *l*_*sq*_). The common language effect size Rho (ρ) is a non-parametric statistic describing the probability that a randomly selected cell from *Lr* will have a greater nAUC than a randomly sampled cell from the ^69^. Thus, ρ can be directly calculated via the Mann-Whitney U-statistic:

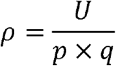

This is also equivalent to the area-under-the-receiver-operating-characteristic-curve (AUC-ROC). Therefore a larger ρ indicates a more accurate prediction result.

We also calculate Cohen’s D as a parametric effect size which provides a measure of robustness and variation in addition to differences between two cell groups:

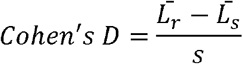

where *s* is the pooled standard deviation from the two groups, i.e., 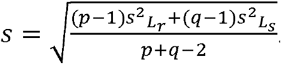. A value of 0.8 or higher is typically viewed as a large effect size and above^37^.

### Cellular drug response prediction via CaDDReS-Sc and Beyondcell

CaDDReS-Sc by default supports only the GDSC drug screen. While the CaDRReS-Sc github repository suggests flexibility to train a new model based on other drug response datasets like CTRP, this pipeline was not applied to CTRP in neither the original manuscript nor github. To allow for comparison between other similar methods, we used CTRPv2. To calculate the weight for each sample-drug pair that is determined by the logistic weight function, we used max concentrations and AUC as the original pipeline learned weights from max concentrations and IC50s. To generate predictions on the single cell samples, we defined the kernel features as the correlations between CCLs and single cell samples. For Beyondcell, we utilized gene signatures generated from only the CTRPv2 database to infer cellular drug sensitivity scores (Beyondcell Scores or BCS, Supplementary Information 1). Since a higher BCS indicates higher sensitivity, we compared sensitive cells against resistant cells to ensure consistent directions with the rest of the methods. Bootstrap aggregation was performed, and performance was summarized across 50 applications of scIDUC and CaDRReS-Sc where 95% of bulk samples were randomly selected for each application. Given that Beyondcell provides pre-trained signatures, we sampled 95% of up- and down-regulated genes without replacement as a bootstrap experiment.

### Cell Culture and Reagents

The DU145 prostate cancer cell line was obtained from American Type Culture Center (ATCC) and cultured in RPMI 1640 medium (Thermo Fisher Scientific), supplemented with 10% fetal bovine serum (FBS) (Gibco, Thermo Fisher Scientific) and maintained at 37 °C with 5% CO_2_. A docetaxel-resistant cell line model for DU145 was established by chronically exposing the parent cell line to stepwise increasing concentrations of docetaxel as previously described^70,71^. Both cell lines were periodically monitored for mycoplasma using the Universal Mycoplasma Detection Kit following the manufacturer’s protocol (ATCC). In vitro drug screening in both the docetaxel-resistant and control DU145 models was performed using either vemurafenib (PLX4032; CAS No. 918504-65-1) or docetaxel (RP-56976; CAS No. 114977-28-5) obtained from MedChem Express (Monmouth Junction, NJ, USA) dissolved in dimethylsulfoxide (DMSO) to obtain stock concentrations of 100mM for vemurafenib or 5mM for docetaxel.

### Drug Screening in Docetaxel-Resistant and Control DU145 Cell Lines

Docetaxel-resistant and control DU145 cells were trypsinized, harvested, counterstained with Hoechst 33342 Fluorescent Stain (Thermo Scientific, Pierce Biotechnology, Rockford, IL) and resuspended in full growth media to 5x10^4^ cells per mL prior to plating in 96-well microplates (Thermo Scientific) using a seeding density of 5x10^3^ cells per well and allowed to attach for 24 hours. Following incubation, cells were treated with different concentrations of either docetaxel ranging from 0.92nM to 6uM, or vemurafenib ranging from 2.5uM to 50uM. Cell viability for each well was measured following a 72-hour drug exposure using the WST-1 assay [(Roche Applied. Science, Penzberg, Upper Bavaria, Germany) following the manufacturer’s protocol. Absorbance at the 450 nm wavelength was assessed using the Synergy HTX Multi-Mode Plate Reader (BioTek, Winooski, VT)]. Absorbance values for each well were used to calculate percent viability relative to the no drug condition. Results are reported as a mean and standard deviation of three independent biological experiments, each containing three technical replicates for each experimental condition.

## Data Availability

We have provided the data and scripts used in this study through DOI 10.17605/OSF.IO/Z9E2X (https://osf.io/z9e2x/).

## Authors’ contributions

Conceptualization, W.Z., D.M., Y.H., and R.S.H.; methodology, W.Z., D.M., and R.S.H.; software, W.Z. and D.M.; experimental validation: A.L., S.J., and I.G.A.; formal analysis, W.Z. and D.M.; prospective analysis: W.Z.; investigation, W.Z., D.M., A.L., R.F.G., Y.H.; writing—original draft preparation, W.Z. and D.M.; writing—review and editing, W.Z., D.M., A.L., R.F.G., Y.H., A.G.P., and R.S.H.; visualization, W.Z.; supervision, A.G.P. and R.S.H.; project administration, W.Z., D.M., and R.S.H.; funding acquisition, W.Z., A.G.P., and R.S.H.

## Acknowledgement

This study was supported by NIH/NCI Grants R01CA204856 (R.S.H.) and the University of Minnesota (UMN) OACA Faculty Research Development grant (R.S.H.). R.S.H. also received support from NIH/NCI R01CA229618, a research grant from the Avon Foundation for Women and the University of Minnesota (UMN) OACA GIA award. W.Z. received the UMN IDF Fellowship.

