## Supplemental Information for "Inferring therapeutic vulnerability within tumors through integration of pan-cancer cell line and single-cell transcriptomic profiles"

**Supplementary Information**

### 1. Brief overview of other single-cell drug response prediction algorithms

We briefly introduce the methodologies of current competing methods. We compared performances of scIDUC against that of Beyondcell and CaDRReS-Sc in our study using a variety of single-cell (SC) datasets with known drug sensitivity information[1,2]. Additionally, Chen et al. have developed a deep learning (DL) approach–scDEAL based on a variational autoencoder–to infer sensitivities to drugs in scRNA-seq data[3]. Similarly, Zheng et al. used an adversarial learning approach and developed SCAD[4]. These DL approaches utilize strictly binarized drug response (sensitive vs. resistant) instead of a continuous value, which better reflects drug response propertities[5]. ASGARD imputes SC drug sensitivities by scanning for drugs that associate with reversed gene expression from disease samples to normal samples[6]. As a result, it requires both disease and normal scRNA-seq data from the same subjects, which is a limiting factor for its utility.

In scIDUC, the CCA integration process involves singular value decomposition (SVD) on the similarity matrix between SC:

$$SVD(X_{bk}X_{sc}{}^{T}) = UDV^{T}$$

Where: $X_{bk} \in \mathbb{R}^{n_1 x g}$ and $X_{sc} \in \mathbb{R}^{n_2 x g}$

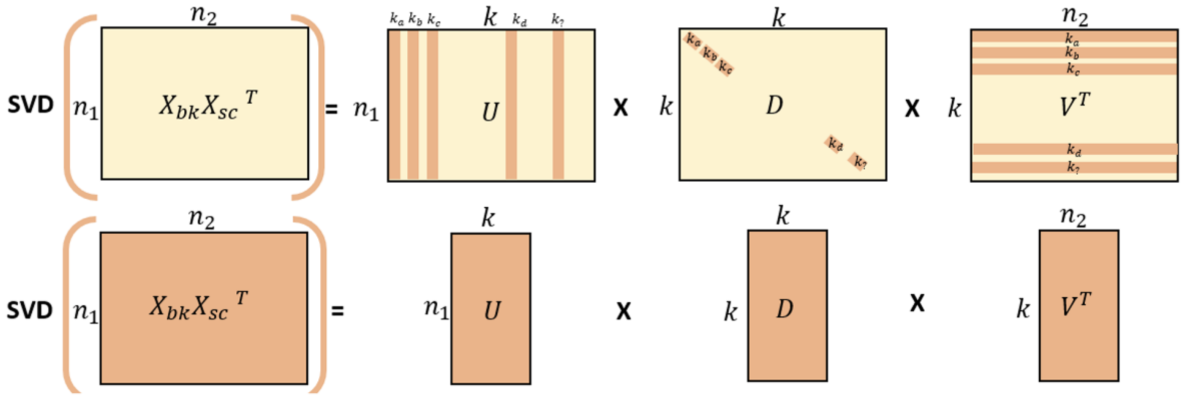


Low rank approximation of $X_{bk}X_{sc}{}^{T}$:

$$Z_{bk} = U\sqrt{D} \in \mathbb{R}^{n_1 x\, k} \qquad Z_{sc} = \sqrt{D}V \in \mathbb{R}^{n_2 x\, k}$$

**Figure S1 Integration of CCL RNA-seq and single-cell RNA-seq datasets via CCA.**

**<u>CaDRReS-Sc:</u>** CaDRReS-Sc is a machine learning framework used for cancer drug response prediction based on single-cell RNA-sequencing data. It extends a previously established method, CaDRReS, calibrated for higher accuracy of drug response prediction based on single cell data. In brief, it is a matrix factorization model, learning a latent pharmacogenomic space that captures the relationship between drug response profiles and transcriptomic data derived from cells, cell clusters, cell lines, or patients. Cell line data screened across a panel of drugs are used for model training to obtain a more robust drug response prediction through sharing information across drugs. The objective function learns the pharmacogenomic space, incorporating a logistic weight function ($C_{iu}$) to assign a weight for each sample-drug pair, reducing noise from extrapolation errors in IC50 values from the dose-response curve fitting step. The objective function that is minimized is defined below.

$$Minimize\ \frac{1}{2}\frac{\Sigma_i \Sigma_u (d_u c_{iu} \left(s_{iu} - \hat{s}_{iu}\right)^2)}{k} + \text{regularization}$$

Where:

$$\hat{s}_{ui} = \mu + b_i^Q + b_u^P + q_i * p_u \text{ which is equivalent to } \hat{s}_{ui} = \mu + b_i^Q + b_u^P + q_i * (X_u * W_p)^T C_{iu} = min(f\left(s_{iu}, o_i, l\right), f\left(\hat{s}_{iu}, o_i, l\right))$$

$$s_{ui} = -\log log(IC50)_2, \text{ The observed sensitivity score of sample } u \text{ for drug } i$$

$$k = \text{The total number of drug sample pairs}$$

$$\mu = \text{The overall mean drug response}$$

$$b_i^Q = \text{The bias for drug } i$$

$$b_u^p = \text{The bias of the unseen sample } u$$

$$q_i \epsilon R^f = \text{Drug } i \text{ in the } f \text{ dimensional latent space}$$

$$p_u \epsilon R^f = \text{Sample } u \text{ in the } f \text{ dimensional latent space}$$

$$W_p \epsilon R^{dxf} = \text{A transformation matrix, projecting transcriptomic kernel features } X_u R^d$$

$$\text{for each sample onto a pharmacogenetic space}$$

**Beyondcell:** Beyondcell requires a scRNAseq expression matrix and a collection of drug signatures to compute a scaled Beyondcell enrichment score (BCS), ranging from 0-1. This score indicates the activity of a signature in the expression matrix or how susceptible each cell is based on the analyzed gene signature where a high and low BCS indicates concordance and discordance between the signature and the analyzed cell, respectively. The BCS is defined in Additional file 1 of the original publication. Alternatively, it is included below.

A signature is obtained from a differential expression analysis and consists of drug perturbation, containing transcriptional changes induced by a drug, or drug sensitivity, reflecting the transcriptional status of sensitivity or resistance prior to drug treatment. Alternatively, the user can provide a GMT file/ranked matrix. If functional signatures are applied, the BCS can be used to evaluate the cell's functional status. Therefore, depending on what collection is used, the BCS can measure the cell perturbation susceptibility or the predicted sensitivity to a given drug. If a gene signature has separate sets of upregulated and downregulated genes and is therefore bidirectional, the BCS is calculated for each signature mode. The individual sum of the expression is calculated and divided by the number of genes in the given signature that are present in the scRNAseq expression matrix.

The BCS is calculated for each drug-cell pair, resulting in a BCS matrix used to determine the presence of therapeutic clusters within the scRNAseq data, visualized using a UMAP. These clusters represent tumor cell subpopulations with distinct shared drug behavior, and the therapeutic differences among the cell populations guide drug selection to nominate cancer-specific treatments. Drug selection is performed using a sensitivity-based ranking, which prioritizes the best drug hits.

For each signature analyzed, a switch point (SP) is calculated, which represents the value in the 0-1 scale where cells switch from down-regulated to up-regulated status. Tumors which are the most therapeutically homogenous will either have a SP of 0, indicating all cells are sensitive to a drug, or 1, indicating all cells are resistant to a drug. A heterogeneous response toward a drug is indicated by a SP between 0-1.

Let $X = (x_{ij})$ be a single-cell expression matrix with $n$ genes contained in $I = \{i_1, \dots, i_n\}$ and $m$ cells contained in $J = \{j_i, \dots, j_m\}$. Also, consider we have a geneset $GS_M = \{S_{M,1}, \dots, S_{M,p}\}$ which consists in a set of $p$ sets of genes (signatures, denoted as $S$ ) per mode in $M = \{UP \vee DN\}$.

To compute the BCS, the following steps are taken:

First, we calculate the intersection between $I$ and each $S_M$ in $GS_M$:

$$G_{M,S_M} = I \cap S_M \quad for\ S_M\ in\ GS_M$$

Second, we obtain the submatrix of $X$ that contains only the genes present in $G_{M,S_M}$. We denote this submatrix as $Y = (y_{gj})$, with $q$ genes contained in $G_{M,S_M} = \{g_1, \dots, g_q\}$. The raw score for each cell $j$ and signature $S_M$ within a mode $M$ is equal to the mean expression of the genes belonging to $G_{M,S_M}$:

$$raw_{M,S_M,j} = \bar{y_j} = \frac{1}{|G_{M,S_M}|} \cdot \sum_{k=1}^{q} y_{kj}$$

Then, the raw scores are normalized, using the mean and the standard deviation of the gene expression.

$$norm_{M,S_M,j} = raw_{M,S_M,j} \cdot f$$

The normalization factor $f$ can be decomposed as follows:

$$f = \frac{\sum_{k=1}^{q} y_{kj} - \sqrt{\frac{\sum_{k=1}^{q}(y_{kj} - \bar{y_j})^2}{q-1}}}{\bar{y_j} + \sqrt{\frac{\sum_{k=1}^{q}(y_{kj} - \bar{y_j})^2}{q-1}}}$$

$$f = \frac{sumexpr - sd}{mean + sd}$$

Thus, the higher the standard deviation the lower $f$. This results in a higher penalization of the raw BCS for cells with outlier genes within $G_{M,S_M}$. Moreover, the lower the $mean$ and $sumexpr$, the lower $f$, which further penalizes cells with a great number of zeros.

Finally, we operate on the normalized scores and scale the results between [0, 1] for each signature $S_M$.

$$BCS_{S_M,j} = \begin{cases} (norm_{UP,S_M,j} - norm_{DN,S_M,j})\ [0,1] & \text{if } M = \{UP, DN\} \\ norm_{UP,S_M,j}\ [0,1] & \text{if } M = \{UP\} \\ -norm_{DN,S_M,j}\ [0,1] & \text{if } M = \{DN\} \end{cases}$$

**scDEAL:** scDEAL integrates a bulk RNA-seq dataset (typically pan-cancer cell line profiles) and a scRNA-seq dataset by minimizing a loss function of maximum mean discrepancy (MMD):

$Loss_{MMD}(E_b(X_b), E_s(X_s)) = |\frac{1}{n}\sum_{i=1}^{n} \phi(x^i{}_b) - \frac{1}{m}\sum_{j=1}^{m} \phi(x_s{}^j)|_H$, where $X_b = \{x_b{}^i\}_{i\in\{1,2,\ldots,n\}}$ is the bulk RNA-seq with $n$ samples and $X_s = \{x_s{}^j\}_{j\in\{1,2,\ldots,m\}}$ is the scRNA-seq with $m$ cells. $\phi$ maps original data into a universal reproducing kernel Hilbert space (RKHS) and $|.|_H$ indicates the RKHS norm measuring distances between two vectors. Such a loss function pursues similar distributions between the two data sources. By incorporating the additional MMD loss into the optimization of an encoder model for predicting drug response, continuous probability scores $Y_s$ are produced for scRNA-seq data. scDEAL then binarizes cellular drug response using a 0.5 probability threshold. The available scDEAL Python package currently only supports the five scRNA-seq experiment datasets appeared in Chen et al.

#### 2. Information of scRNA-seq data used in the paper

Basic properties of the scRNA-seq used in this manuscript is shown in Table S1. We have preprocessed each dataset using the scanpy package. For each one we included cells with at least 200 RNAs detected and genes detected in at least 3 cells. Cells with high percentages of mitochondria genes were filtered out. We did not select any high variability genes as scIDUC utilizes drug response relevant genes (DRGs) in the input datasets.

**Table S1 Information of the scRNA-seq datasets**

| Data Name | Authors | No. of Cells | Sample Source | Disease | Drug Name | Availability |
|---|---|---|---|---|---|---|
| Lung-PC9 | Kong et al. | 507 | PC9 cell line | Lung cancer | Gefitinib | GSE112274 |
| AML-PDX | Bell et al. | 1472 | MLL-AF9 | Acute Myeloid Leukemia | I-BET-151 | GSE110894 |
| Breast-MCF7 | Ben-David et al.[a] | 2899 | MCF7 cell line | Breast cancer | Bortezomib | GSE114462 |
| RMS-oPDX | Patel et al. | 5643 | Patient-derived xenografts | Rhabdomyosarcoma (RMS) | SN-38 and EGFRis | GSE174376[b] |
| CRPC-CCLs | Schnepp et al. | 324 | PC3 and DU145 cell lines | Castration-resistant prostate cancer (CRPC) | Docetaxel | GSE140440 |
| PDAC-CFPAC1 | Raghavan et al. | 2042 | CFPAC1 cell line | Pancreatic ductal adenocarcinoma (PDAC) | SN-38 and Paclitaxel | Single Cell Portal #1644[c] |

[a] For the Ben-David dataset, we treated t0 cells as Bortezomib sensitive cells and t96 cells as resistant cells.

[b] The RMS data in our study was obtained directly from Dr. Anand G. Patel.

[c] CFPAC1 cell line data presented in Figure 4 in Raghavan et al. (2021) was used in our paper.

3. Performances of scIDUC and other competing methods

**Table S2 scIDUC performance with different integration algorithms and different cell-to-DRG ratios.**

| Data | Method | Metric | SC-DRG Ratio | | | | | | |
|---|---|---|---|---|---|---|---|---|---|
| | | | **10** | **5** | **2** | **1** | **0.5** | **0.3** | **0.2** |
| AML-PDX | CCA Integration | -LOG10(P-Value) | 15.11 (13.23) | 23.23 (19.39) | 49.56 (46.08) | 53.23 (43.31) | 46.6 (40.16) | 41.86 (38.96) | 58.24 (56.72) |
| AML-PDX | CCA Integration | Cohen's D | 0.61 (0) | 0.75 (0) | 1.17 (0) | 1.17 (0.01) | 1.11 (0) | 1.06 (0.02) | 1.31 (0.02) |
| AML-PDX | CCA Integration | Rho | 0.66 (0) | 0.69 (0) | 0.8 (0) | 0.8 (0) | 0.79 (0) | 0.78 (0) | 0.83 (0) |
| AML-PDX | NMF Integration | -LOG10(P-Value) | 0.55 (0) | 1.6 (0) | 0.33 (0) | 0.96 (1.34) | 4.58 (4.72) | 6.93 (7.77) | 6.72 (5.96) |
| AML-PDX | NMF Integration | Cohen's D | 0.27 (0) | 0.06 (0) | 0.26 (0) | 0.37 (0.02) | 1.14 (0.02) | 1.11 (0.02) | 0.87 (0.05) |
| AML-PDX | NMF Integration | Rho | 0.6 (0) | 0.52 (0) | 0.58 (0) | 0.64 (0.01) | 0.8 (0) | 0.8 (0) | 0.74 (0.01) |
| AML-PDX | No Integration | -LOG10(P-Value) | 15.11 (13.23) | 23.23 (19.39) | 49.56 (46.08) | 53.23 (43.31) | 46.6 (40.16) | 41.86 (38.96) | 58.24 (56.72) |
| AML-PDX | No Integration | Cohen's D | 0.12 (0.06) | -0.2 (0.06) | 0.12 (0.16) | 0.34 (0.06) | 0.26 (0.06) | -0.16 (0.06) | -0.1 (0.07) |
| AML-PDX | No Integration | Rho | 0.54 (0.02) | 0.45 (0.02) | 0.53 (0.05) | 0.59 (0.02) | 0.57 (0.02) | 0.46 (0.02) | 0.48 (0.02) |
| Breast-MCF7 | CCA Integration | -LOG10(P-Value) | 6.98 (3.75) | 26.78 (15.94) | 55 (56.14) | 41.5 (33.58) | 72.17 (54.36) | 106.57 (103.98) | 50.39 (100.46) |
| Breast-MCF7 | CCA Integration | Cohen's D | 0.24 (0.01) | 0.39 (0.01) | 0.11 (0.01) | 0.71 (0.04) | 0.93 (0.05) | 1.24 (0.2) | 1.79 (0.14) |
| Breast-MCF7 | CCA Integration | Rho | 0.58 (0) | 0.62 (0) | 0.53 (0) | 0.7 (0.01) | 0.76 (0.01) | 0.82 (0.04) | 0.9 (0.02) |
| Breast-MCF7 | NMF Integration | -LOG10(P-Value) | 3.33 (0) | 0.28 (0) | 9.61 (12.26) | 5.99 (6.41) | 12.65 (13.8) | 11.72 (13.09) | 16.16 (19.22) |
| Breast-MCF7 | NMF Integration | Cohen's D | 0.06 (0) | 0.25 (0) | -0.41 (0.04) | 0.05 (0.05) | 0.88 (0.16) | 0.58 (0.21) | 1.27 (0.31) |
| Breast-MCF7 | NMF Integration | Rho | 0.52 (0) | 0.57 (0) | 0.39 (0.01) | 0.51 (0.02) | 0.74 (0.04) | 0.68 (0.06) | 0.86 (0.05) |
| Breast-MCF7 | No Integration | -LOG10(P-Value) | 6.98 (3.75) | 26.78 (15.94) | 55 (56.14) | 41.5 (33.58) | 72.17 (54.36) | 106.57 (103.98) | 50.39 (100.46) |
| Breast-MCF7 | No Integration | Cohen's D | 0.13 (0.09) | -0.4 (0.2) | -0.87 (0.12) | -0.21 (0.1) | -0.34 (0.1) | -0.23 (0.12) | -0.23 (0.12) |
| Breast-MCF7 | No Integration | Rho | 0.54 (0.02) | 0.39 (0.05) | 0.27 (0.03) | 0.44 (0.03) | 0.41 (0.03) | 0.43 (0.03) | 0.44 (0.03) |
| Lung-PC9 | CCA Integration | -LOG10(P-Value) | 13.58 (12.84) | 14.62 (13.19) | 22.71 (14.62) | 21.9 (18.97) | 14.26 (8.64) | 22.27 (11.38) | 28.71 (14.94) |
| Lung-PC9 | CCA Integration | Cohen's D | 2.02 (0) | 2.15 (0.18) | 2.75 (0) | 2.89 (0.01) | 2.06 (0.09) | 2.58 (0.05) | 3.01 (0.11) |
| Lung-PC9 | CCA Integration | Rho | 0.92 (0) | 0.93 (0.01) | 0.96 (0) | 0.96 (0) | 0.91 (0.01) | 0.95 (0) | 0.97 (0) |
| Lung-PC9 | NMF Integration | -LOG10(P-Value) | 1.68 (0) | 2.92 (0) | 24.71 (0) | 3.88 (0) | 0.81 (0.89) | 0.96 (1.03) | 0.74 (0.59) |
| Lung-PC9 | NMF Integration | Cohen's D | 0.96 (0) | 1.68 (0) | 1.26 (0) | 1.28 (0) | 1.61 (0.09) | 1.71 (0.1) | 2.01 (0.08) |
| Lung-PC9 | NMF Integration | Rho | 0.85 (0) | 0.89 (0) | 0.82 (0) | 0.83 (0) | 0.9 (0.01) | 0.9 (0.01) | 0.93 (0.01) |
| Lung-PC9 | No Integration | -LOG10(P-Value) | 13.58 (12.84) | 14.62 (13.19) | 22.71 (14.62) | 21.9 (18.97) | 14.26 (8.64) | 22.27 (11.38) | 28.71 (14.94) |
| Lung-PC9 | No Integration | Cohen's D | -0.19 (0.13) | -0.34 (0.1) | -0.97 (0.1) | -0.52 (0.22) | -0.86 (0.16) | -1.09 (0.12) | -1.2 (0.16) |
| Lung-PC9 | No Integration | Rho | 0.43 (0.04) | 0.37 (0.03) | 0.25 (0.02) | 0.36 (0.06) | 0.27 (0.04) | 0.22 (0.02) | 0.2 (0.03) |

[a] values are presented as mean (SD).

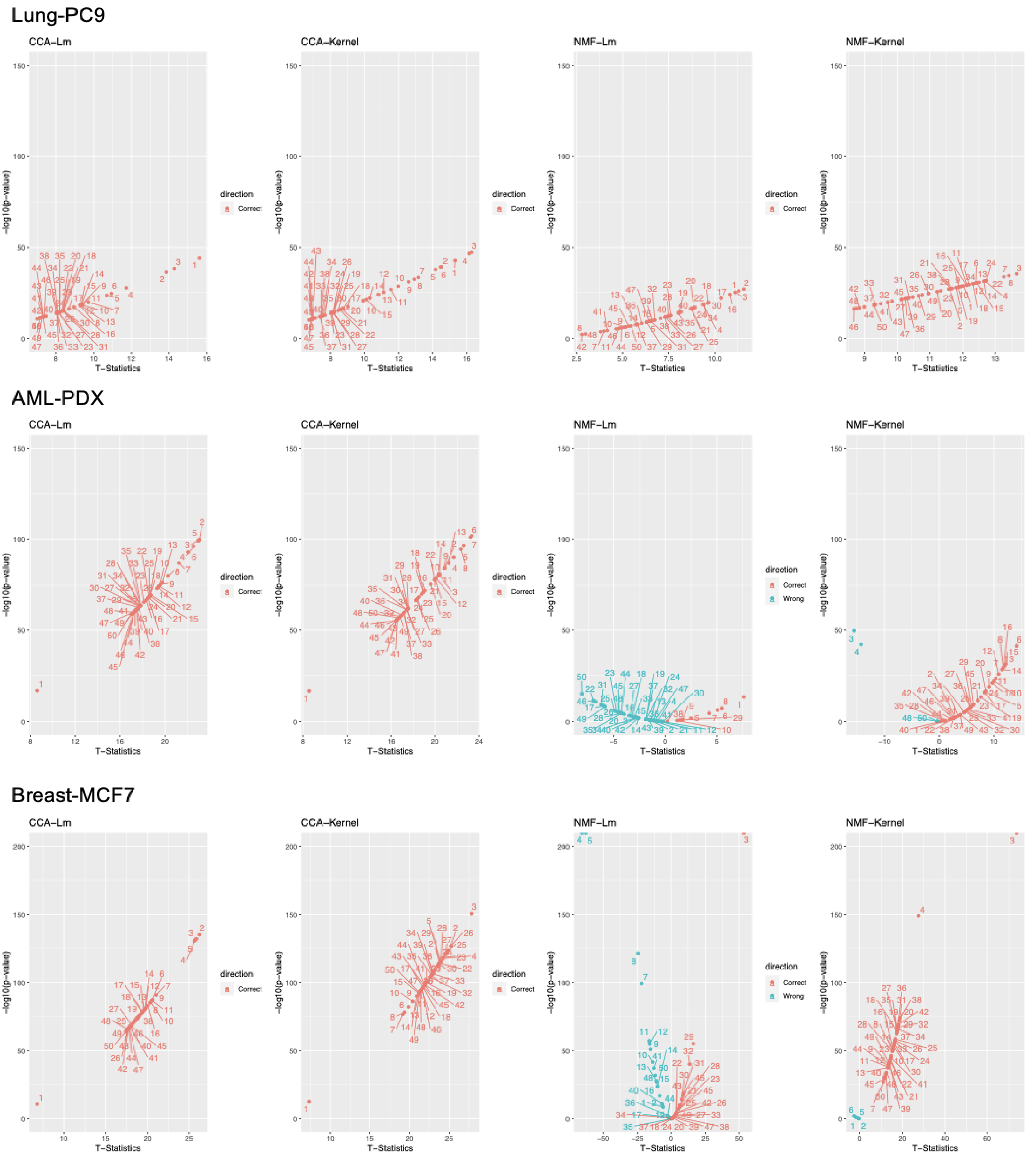


**Figure S2 Robustness of CCA and NMF based integration for single-cell drug response prediction.** Combinations of integration methods (CCA or NMF) and drug response models (Lm: linear model;

Kernel: kernelized nonparametric regression) were tested. Inner dimensions of CCA or NMF ranging from 1 to 50 were tested for robustness. T-tests results comparing predicted nAUC between true resistant and sensitive groups are shown; a t-statistic larger than 0 indicates correct direction.

#### Table S3 Benchmarking scIDUC against other competing methods[a].

| Data | Method | Cohen's D | -LOG10(P-Value) | Rho |
|---|---|---|---|---|
| CRPC-CCLs | Beyondcell | 0.94 (0.02) | 14.32 (0.6) | 0.74 (0) |
| CRPC-CCLs | CaDRReS-Sc | 0.29 (0.02) | 2.01 (0.17) | 0.57 (0) |
| CRPC-CCLs | scIDUC (SC-DRG=0.2) | 1.52 (0.07) | 33.25 (2.28) | 0.87 (0.01) |
| CRPC-CCLs | scIDUC (SC-DRG=0.9) | 1.78 (0.12) | 42.11 (4.07) | 0.89 (0.02) |
| PDAC-CFPAC1 | Beyondcell | 0.55 (0) | 33.37 (0) | 0.65 (0) |
| PDAC-CFPAC1 | CaDRReS-Sc | 0.39 (0.01) | 17.36 (0.71) | 0.39 (0) |
| PDAC-CFPAC1 | scIDUC (SC-DRG=0.2) | 1.2 (0.11) | 138.46 (22.31) | 0.81 (0.02) |
| PDAC-CFPAC1 | scIDUC (SC-DRG=0.9) | 1.22 (0.25) | 143.98 (44.6) | 0.8 (0.05) |
| RMS-oPDX | Beyondcell | 0.94 (0.02) | 14.32 (0.6) | 0.74 (0) |
| RMS-oPDX | CaDRReS-Sc | 0.12 (0.01) | 2.66 (0.21) | 0.47 (0) |
| RMS-oPDX | scIDUC (SC-DRG=0.2) | 1.65 (0.17) | 300 (0) | 0.88 (0.02) |
| RMS-oPDX | scIDUC (SC-DRG=0.9) | 2.07 (0.16) | 300 (0) | 0.93 (0.01) |
| AML-PDX | Beyondcell | 0.55 (0) | 33.37 (0) | 0.65 (0) |
| AML-PDX | CaDRReS-Sc | 0.39 (0.01) | 17.36 (0.71) | 0.39 (0) |
| AML-PDX | scIDUC (SC-DRG=0.2) | 1.2 (0.11) | 138.46 (22.31) | 0.81 (0.02) |
| AML-PDX | scIDUC (SC-DRG=0.9) | 1.22 (0.25) | 143.98 (44.6) | 0.8 (0.05) |
| Breast-MCF7 | Beyondcell | 1.37 (0.07) | 238.46 (19.84) | 0.83 (0.01) |
| Breast-MCF7 | CaDRReS-Sc | 1.66 (0.01) | Inf (NaN) | 0.88 (0) |
| Breast-MCF7 | scIDUC (SC-DRG=0.2) | 1.77 (0.15) | 300 (0) | 0.9 (0.02) |
| Breast-MCF7 | scIDUC (SC-DRG=0.9) | 0.77 (0.04) | 300 (0) | 0.72 (0.01) |
| Lung-PC9 | Beyondcell | 2.05 (0.05) | 15.33 (0.48) | 0.92 (0) |
| Lung-PC9 | CaDRReS-Sc | 0.46 (0.06) | 23.04 (10.79) | 0.36 (0.03) |
| Lung-PC9 | scIDUC (SC-DRG=0.2) | 2.94 (0) | 41.47 (0) | 0.97 (0) |
| Lung-PC9 | scIDUC (SC-DRG=0.9) | 2.94 (0) | 41.47 (0) | 0.97 (0) |

[a] The values are presented as mean (SD).

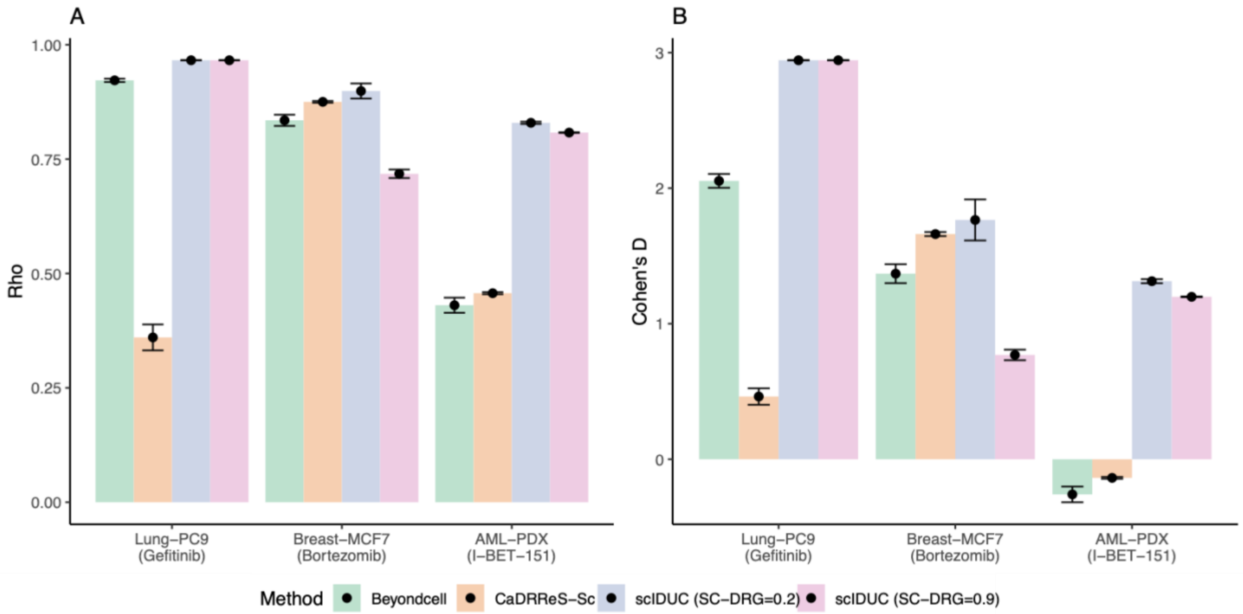


**Figure S3 scIDUC outperforms other methods across three additional scRNA-seq datasets.** For each method, 50 bootstrap samples were generated (see Methods). In all three datasets, scIDUC shows higher Common-language effect Rho (A) and Cohen' D (B) comparing predicted cell response between the true resistant vs. sensitive cell groups than that of other methods (CaDRReS-Sc or Beyondcell).

#### 4. Prospective analysis to capture therapeutic vulnerabilities in PDAC-CFPAC1 cells shaped by TMEs

A

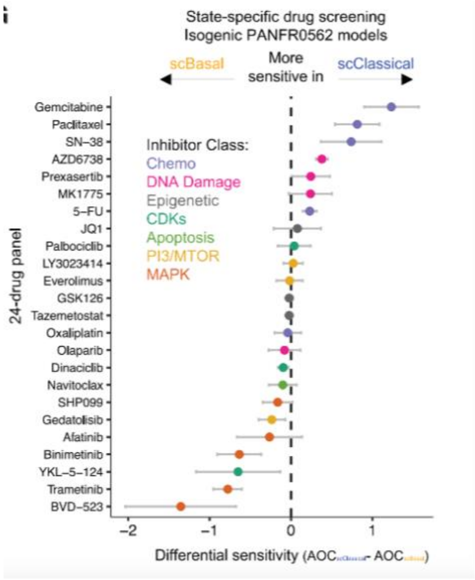


B

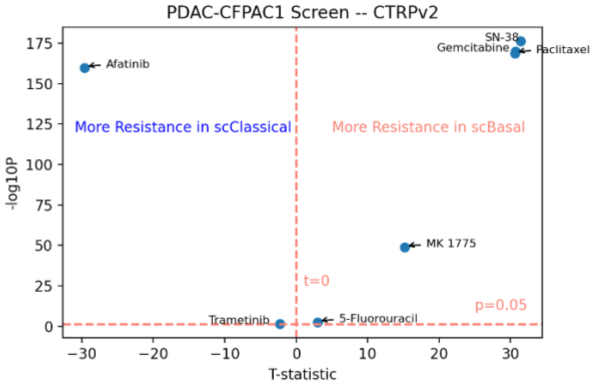


**Figure S4 A.** Drug panel screens showing differential efficacy in a pancreatic ductal carcinoma patient-derived xenograft model with two different subtypes due to different TMEs. This figure was originally generated by Raghavan et al. (Figure 6G in DOI:https://doi.org/10.1016/j.cell.2021.11.017). **B.** Predicted cellular response to various drugs using scIDUC. T-tests results comparing predicted nAUCs between scBasal and scClassical cells are shown.

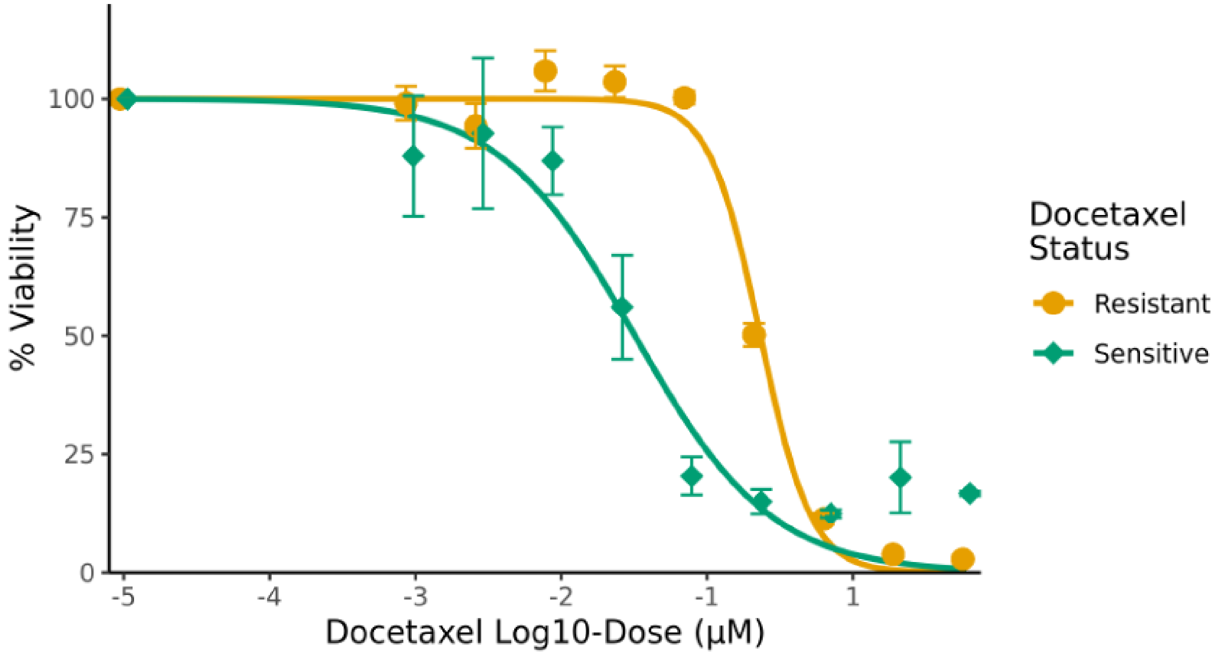

**Figure S5 WST assay showing differential sensitivity to docetaxel among DU145 cells.** Docetaxel-resistant DU145 cells show higher resistance to docetaxel compared to their docetaxel-sensitive counterparts in vitro (two-way ANOVA p<0.0001). At each concentration, mean percent viability ± standard deviation is plotted.

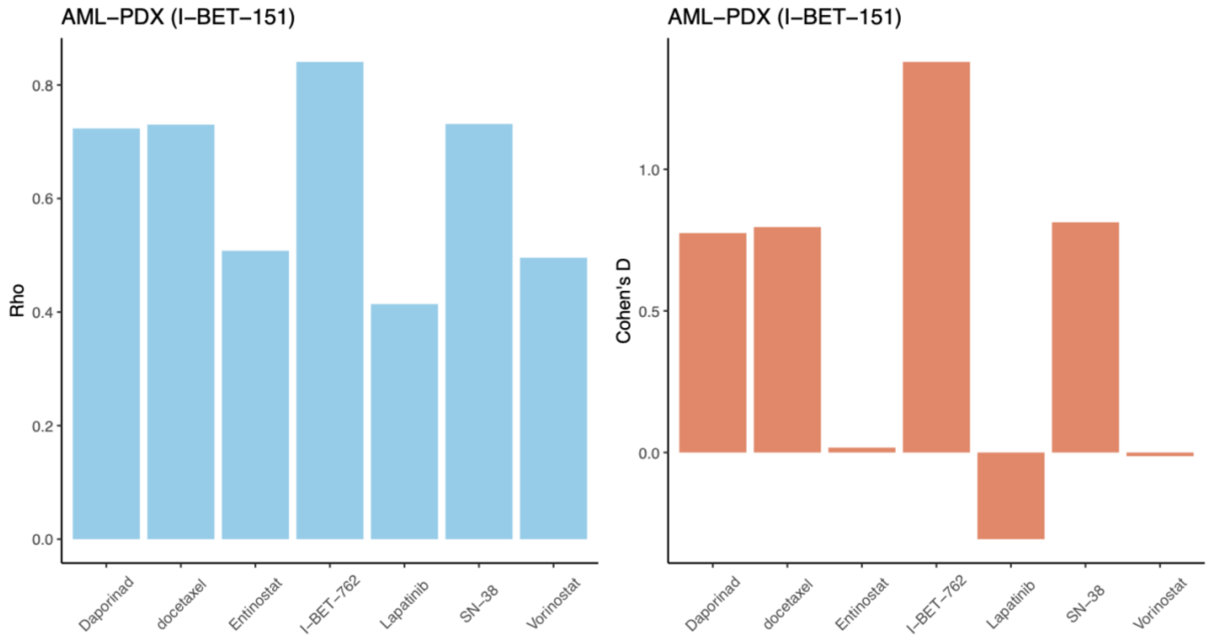


**Figure S6 scIDUC predicts accurate results in the presence of potential bath effects.** Left: Rho statistics comparing predicted nAUCs between I-BET sensitive cells and resistant cells over seven drugs. Right: Cohen's D comparing predicted nAUCs between I-BET sensitive cells and resistant cells over seven drugs.
